# Dual-Method Immune Deconvolution Reveals Subtype-Specific PD-L1 Drivers and a Robust γδ T Cell–M2 Macrophage Axis in Breast Cancer

**DOI:** 10.64898/2026.05.30.728960

**Authors:** Diya Jain, Hari S. Misra

**Author notes:** Corresponding author: Prof. Hari S Misra.

## Abstract

Most computational studies of the breast cancer immune microenvironment rely on a single deconvolution algorithm. Since different tools analyze different parameters, integrated information across a dataset can be missed. Here, we employed two complementary approaches, xCell and a CIBERSORT-style ssGSEA using LM22 signatures, on 1,099 PAM50-classified primary tumors from the TCGA-BRCA cohort. PD-L1 was highest in basal-like tumors (Kruskal–Wallis p < 0.001) and correlated with CD8+ T cells (ρ = 0.65) and M1 macrophages (ρ = 0.67) in that subtype, which fits the standard model of IFN-γ-driven adaptive upregulation. In HER2-enriched cancers, PD-L1 tracked with both effector and regulatory populations simultaneously, while luminal tumors were largely immune-quiet. The most consequential finding involved γδ T cells and M2 macrophages: xCell showed a non-significant correlation (ρ = 0.048, p = 0.11), whereas CIBERSORT-ssGSEA, using curated γδ gene signatures, produced a significant correlation (ρ= 0.565, p < 2.2 × 10⁻¹⁶) that held across all five subtypes. A multivariate model explained 49% of PD-L1 variance (adjusted R² = 0.49), and Cox regression incorporating immune features gave a concordance of 0.60. These results suggest a baseline γδ–M2 immunosuppressive circuit in breast cancer, particularly in ER+ disease, that could be useful to set a therapeutic target.

## 1. Introduction

Breast cancer is a disease characterized by several molecular sub-types. The PAM50 subtypes, basal-like, HER2-enriched, luminal A, luminal B, and normal-like, differ in their transcriptional programs, clinical trajectories, and, importantly, in the immune cells they attract and tolerate (1, 2). These differences are critical for treatment. Immune checkpoint blockade has changed outcomes in several solid tumors; however, in breast cancer its benefits have been mostly confined to triple-negative disease, where pembrolizumab plus chemotherapy improves survival in PD-L1-positive patients (3). For luminal cancers, which constitute the majority of diagnoses, immunotherapy has not been very successful, and further study is needed to understand luminal cancer treatment beyond PD-L1. Mechanistically, PD-L1 reflects an inflamed microenvironment driven by IFN-γ from activated T cells (6). In basal-like breast cancer, PD-L1 tracks with cytotoxic lymphocyte infiltration, consistent with the success of PD-L1-directed therapy in this setting. In other subtypes, the relationship is less clear. HER2-enriched tumors often show intermediate PD-L1 expression alongside a mix of effector and suppressive immune cells. Luminal cancers, though immunologically relatively “quiet”, still harbor immunosuppressive populations, including Tregs, M2-polarised macrophages, and γδ T cells (4, 5, 7). Tumor-associated macrophages (TAMs) and γδ T cells are central to this immunosuppression. TAMs account for nearly half of all immune cells in breast tumor tissue, and their polarization state shapes the tumor immune microenvironment. M1-polarised macrophages present antigen and promote inflammation. M2-polarised macrophages do the opposite: they produce IL-10 and TGF-β, recruit Tregs, remodel the extracellular matrix, and can express checkpoint ligands themselves (8, 9). What determines macrophage polarization in different breast cancer contexts is still understudied. γδ T cells are also becoming increasingly important. They recognize stressed and transformed cells without MHC restriction, which means they can respond to tumors that have evaded classical antigen presentation (10). In TNBC, γδ T cells act as cytotoxic effectors and associate with better outcomes (11). But they are functionally plastic: IL-17-producing subsets can instead promote immunosuppression, angiogenesis, and macrophage reprogramming toward an M2 state via NF-κB signaling (12). Petroni and colleagues (2025) showed that CDK4/6 inhibition in HR+ breast cancer triggers recruitment of IL-17A-secreting γδ T cells, which reprogram TAMs into a CX3CR1+ immunosuppressive state and drive treatment resistance (4). These findings raise an obvious question: is this γδ–macrophage circuit already present at baseline?

Answering this question with greater confidence requires more than one computational method. xCell provides enrichment scores for over 60 cell types and includes a spillover compensation step to reduce false positives from shared markers, but this same step can dampen real signals from rare populations such as γδ T cells (13, 15). CIBERSORT-based approaches use the LM22 signature matrix and detect lymphocyte subsets with higher specificity (16). Applying both approaches together is worth considering because it produces more reliable information than either alone. We applied both methods to the TCGA-BRCA cohort (n = 1,099) with five goals: (i) compare PD-L1 and immune infiltration across all five subtypes; (ii) map subtype-specific PD-L1–immune correlation patterns; (iii) test whether a baseline γδ T cell–M2 macrophage association exists, with cross-method validation; (iv) identify independent predictors of PD-L1 through multivariate modelling; and (v) assess the prognostic value of immune populations. The results produced from this study are consistent across methods and can be used for further validation.

## 2. Results and Discussion

### 2.1 PD-L1 expression varies across molecular subtypes

The PD-L1 (CD274) expression profile across different subtypes of breast cancer was analyzed in silico. PD-L1 expression differed across subtypes (Kruskal–Wallis p = 4.22 × 10⁻⁹). HER2-enriched tumors had the highest median expression (log2CPM = 2.81), followed by basal-like (2.60), normal-like (2.57), luminal A (2.14), and luminal B (2.05) (Figure 1). The higher levels in immunologically active subtypes are consistent with IFN-γ-driven upregulation by infiltrating T cells, while the low levels in luminal cancers match their generally quiescent immune environment. The higher PD-L1 in HER2-enriched tumors compared to basal-like differs from reports that place TNBC/basal at the top. This is likely because PAM50 subtypes do not perfectly overlap with clinical receptor status, and the HER2-enriched class captures a transcriptionally distinct group with its own immune profile. Regardless, the HER2 → basal → luminal gradient confirms what we set out to test: PD-L1 expression tracks with immune activity across the breast cancer subtype spectrum.

**Figure 1.**
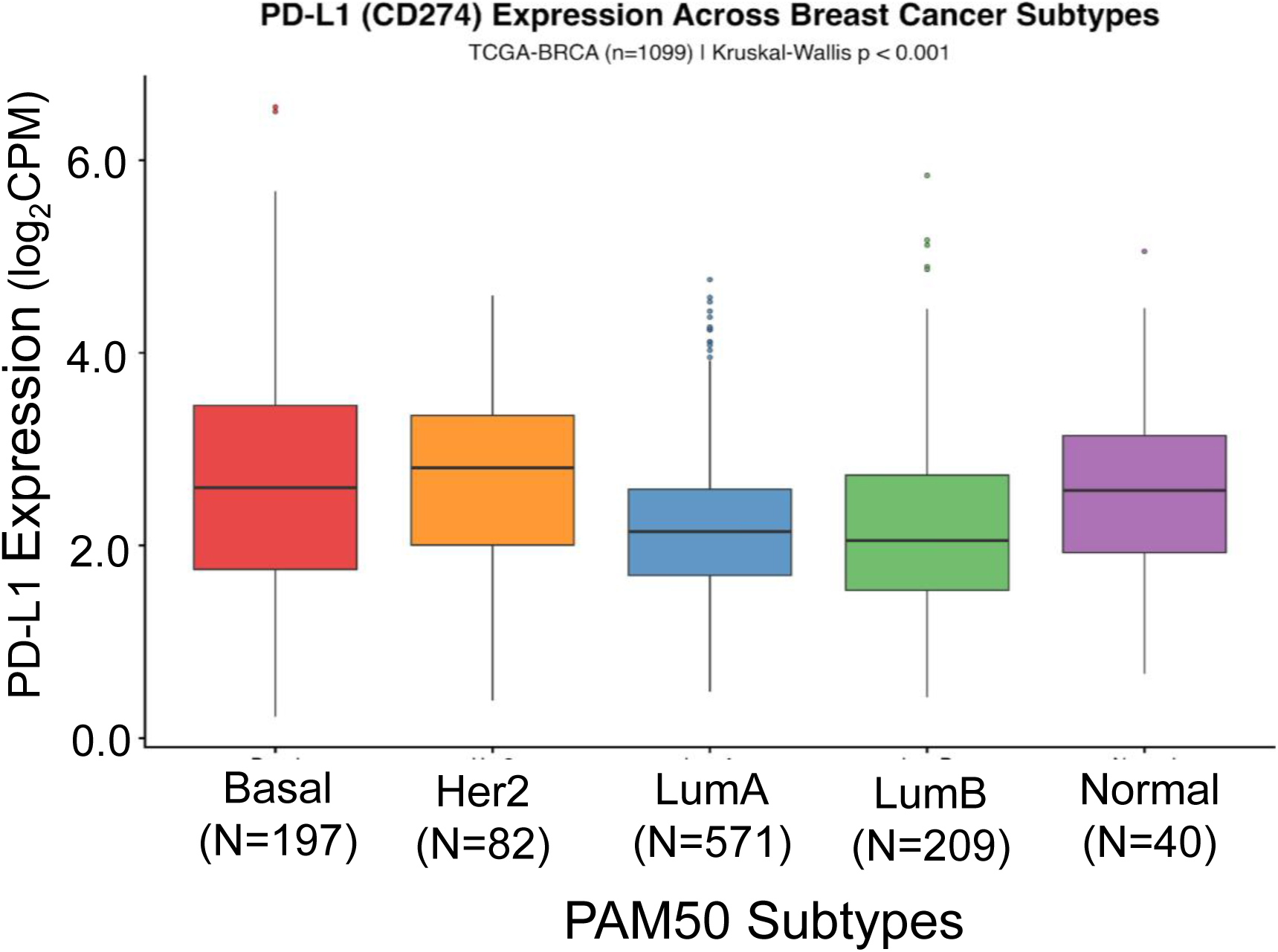
PD-L1 (CD274) expression across breast cancer subtypes. Boxplots of log2CPM in TCGA-BRCA (n = 1,099) stratified by PAM50 subtype. Kruskal–Wallis p = 4.22 × 10⁻⁹.

### 2.2 Immune infiltration patterns are subtype-dependent

Immune infiltration in each breast cancer subtype was analyzed using xCell and CIBERSORT-ssGSEA. The head-to-head comparison for CD8+ T cells and M1 macrophages across both methods is shown in Figure 2. xCell deconvolution revealed clear differences between subtypes. Basal-like tumors had the highest CD8+ T cell and M1 macrophage enrichment, consistent with a cytotoxically active microenvironment. Luminal B and normal-like tumors were enriched for M2 macrophages instead, and Tregs accumulated preferentially in luminal subtypes. NK cell infiltration was uniformly low, while γδ T cell scores were modest but detectable and tended higher in HER2-enriched and luminal B tumors (Figure S1). CIBERSORT-ssGSEA agreed with the xCell analysis. CD8+ T cell and M1 macrophage signatures were again highest in basal-like tumors, and M2 macrophage and Treg signatures dominated the luminal subtypes. However, unlike xCell, which showed modest scores, the CIBERSORT approach picked up notably stronger γδ T cell signals with clearer separation between subtypes (Figure S2). Cross-method correlations were strong for CD8+ T cells, M1 and M2 macrophages, and Tregs, confirming that both methods agree on the broad immune architecture (Figure S3). All Kruskal–Wallis tests for immune cell differences were significant (FDR < 0.001). Taken together, the two deconvolution methods converge on the same subtype-dependent immune infiltration pattern, lending confidence that the picture drawn here is not an artefact of any one algorithm.

**Figure 2.**
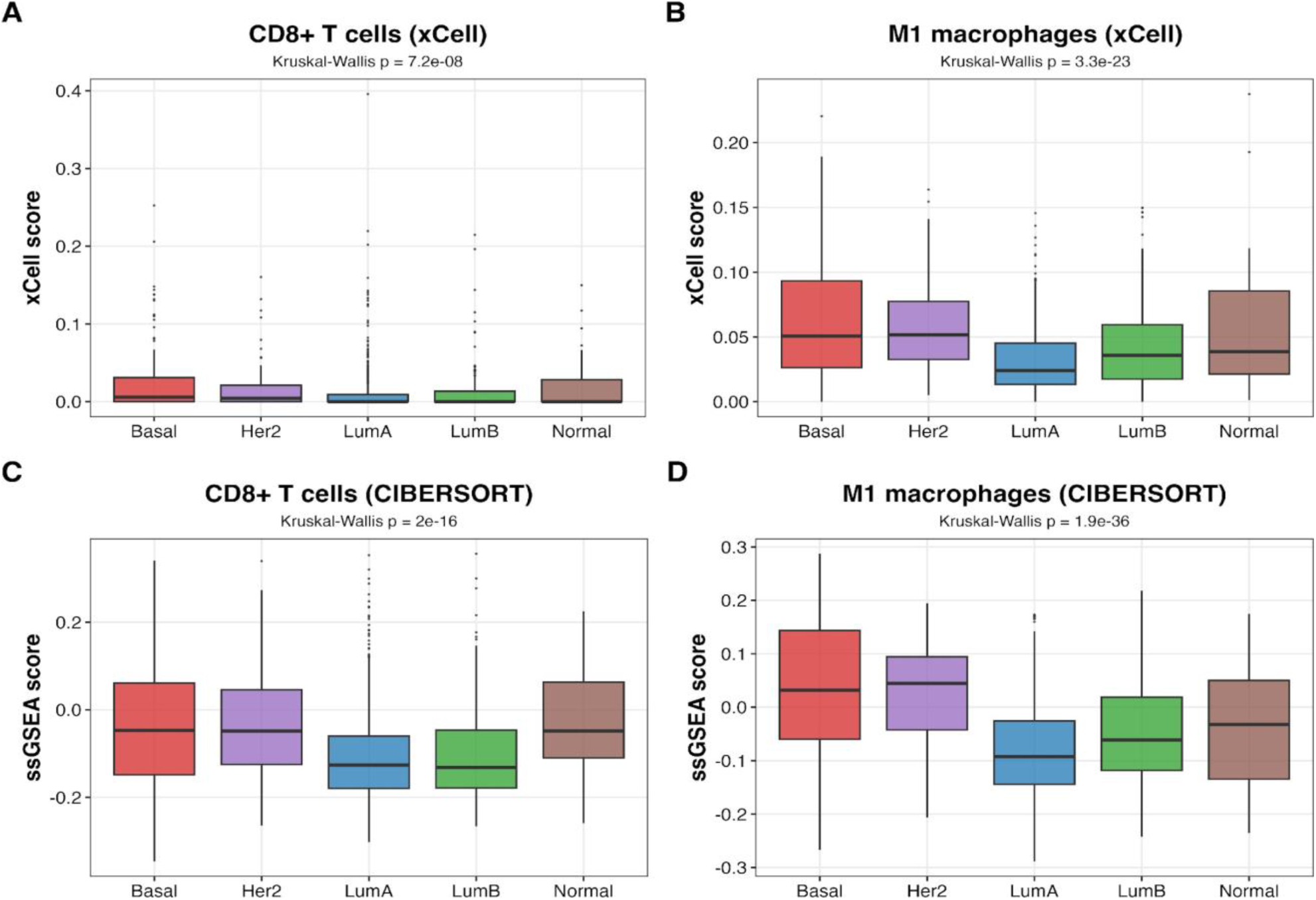
Method-matched comparison of hot-tumour infiltrates. (A) CD8+ T cells by xCell, (B) M1 macrophages by xCell, (C) CD8+ T cells by CIBERSORT-ssGSEA, (D) M1 macrophages by CIBERSORT-ssGSEA. All Kruskal–Wallis FDR < 0.001.

### 2.3 PD-L1 correlates with different immune populations depending on subtype

The association of PD-L1 with individual immune populations across subtypes was next examined. Globally, PD-L1 showed a strong correlation with M1 macrophages (ρ = 0.50), CD8+ T cells (ρ = 0.43), and Tregs (ρ = 0.38), with a weaker M2 macrophage association (ρ = 0.19) and no significant link to NK cells or γδ T cells (Fig. 3A). In basal-like tumors, PD-L1 correlated strongly with M1 macrophages (ρ = 0.67) and CD8+ T cells (ρ = 0.65). In HER2-enriched tumors, PD-L1 correlated with M1 macrophages (ρ = 0.63) but also with Tregs (ρ = 0.43) and γδ T cells (ρ = 0.33), with a weaker CD8 link (ρ = 0.26). Luminal B tumors showed moderate correlations, suggesting that the small fraction of luminal tumors with any immune activity can upregulate PD-L1. Luminal A tumors showed weak or absent correlations across the board (Figure 3B–C).

**Figure 3.**
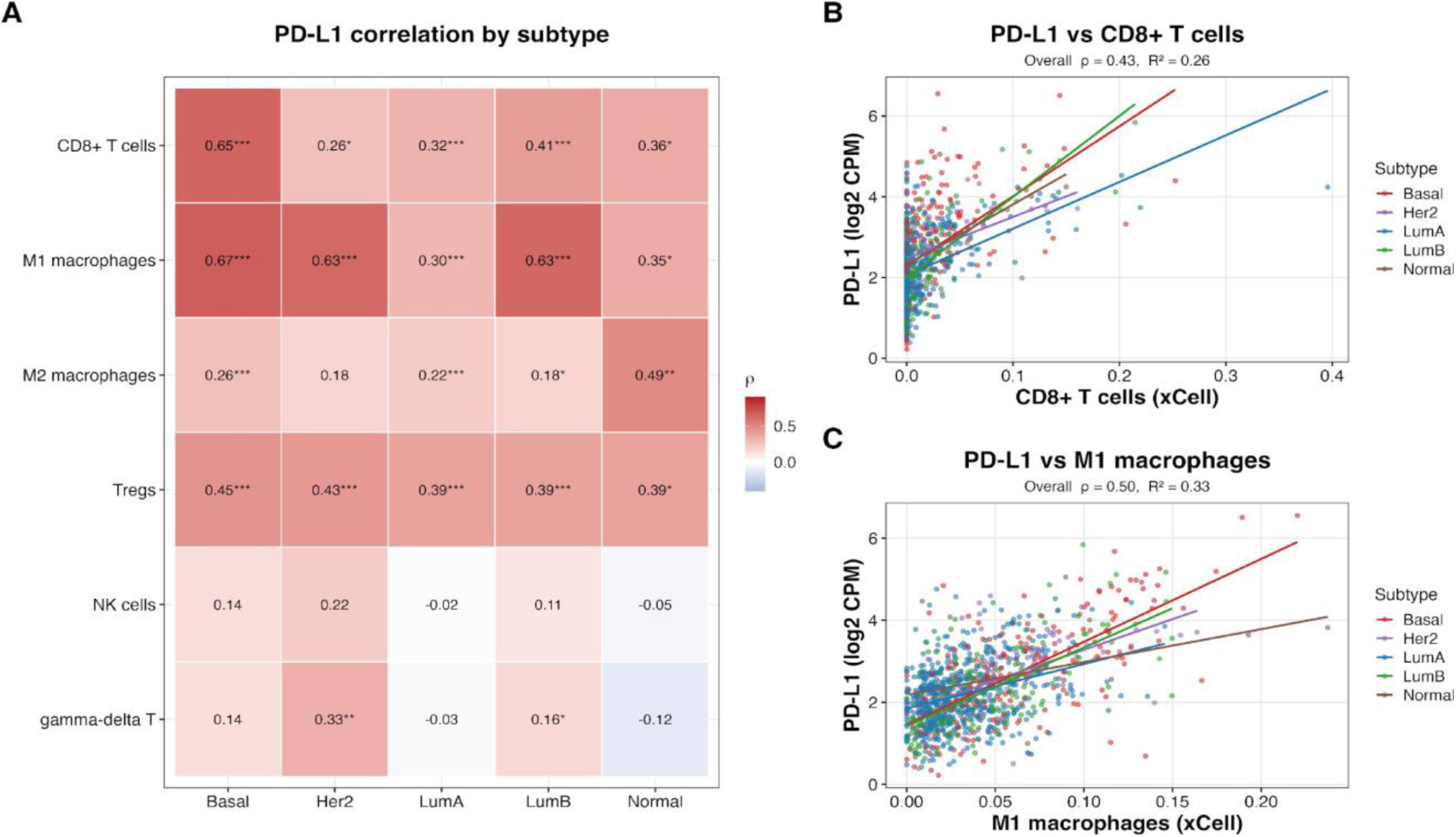
PD-L1–immune correlation structure. Spearman ρ between PD-L1 and six immune cell populations by PAM50 subtype; cell-wise FDR significance shown (*p < 0.05, **p < 0.01, ***p <0.001) (A). Similarly, PD-L1 vs CD8+ T cells and (C) PD-L1 vs M1 macrophages, coloured by subtype; overall ρ and R² from a linear fit are shown in each panel (B).

PD-L1 therefore tracks an active cytotoxic response in basal tumors and may work as a biomarker in that setting. Since PD-L1 testing is already used to determine pembrolizumab eligibility in TNBC (3), this result has direct clinical implications in basal tumors. However, extending PD-L1-based selection to HER2-enriched or luminal subtypes may not be effective, because the same marker co-occurs with both immune activation and immune suppression in these tumors. A patient with high PD-L1 in a HER2-enriched tumor might have an active anti-tumor response, Treg-mediated suppression, or both. Composite biomarkers that integrate subtype, immune cell composition, and checkpoint expression would therefore be a better choice for selecting patients likely to benefit from checkpoint blockade, returning to the central point that PD-L1 does not carry the same meaning in every subtype.

### 2.4 Method choice changes the γδ T cell–M2 macrophage signal

The γδ T cell–M2 macrophage interaction was studied using both xCell and CIBERSORT-ssGSEA, and the different results obtained from the two approaches illustrate why running multiple methods matters in in silico immune profiling (Figure 4). The correlation between γδ T cells and M2 macrophages across all 1,099 tumors was positive but non-significant with xCell (ρ = 0.048, p = 0.11) (Figure 4A). Using CIBERSORT-ssGSEA, a strong positive correlation (ρ = 0.565, p < 2.2 × 10⁻¹⁶; Fig 4B) was observed between the curated γδ T cell gene signature (TRGC1, TRGC2, TRDC, TRDV2, TRGV9) and M2 macrophages. This association was not confined to one subtype: it held across all five, with Basal (ρ = 0.58), Lum A (ρ = 0.52), Lum B (ρ = 0.59), HER2 (ρ = 0.53), and Normal (ρ = 0.44) of tumors. An R² of approximately 0.32 for the CIBERSORT fit gives an effect-size anchor: roughly a third of the γδ variance in tumors is shared with M2 macrophage variance. The discrepancy between methods likely reflects xCell’s spillover compensation, which was designed to reduce false positives from genes shared across cell types and works well for most populations. For rare cells like γδ T cells, which have a limited transcriptomic footprint, the compensation appears to attenuate genuine signals. Direct ssGSEA scoring of specific gene sets avoids this and captures the association.

**Figure 4.**
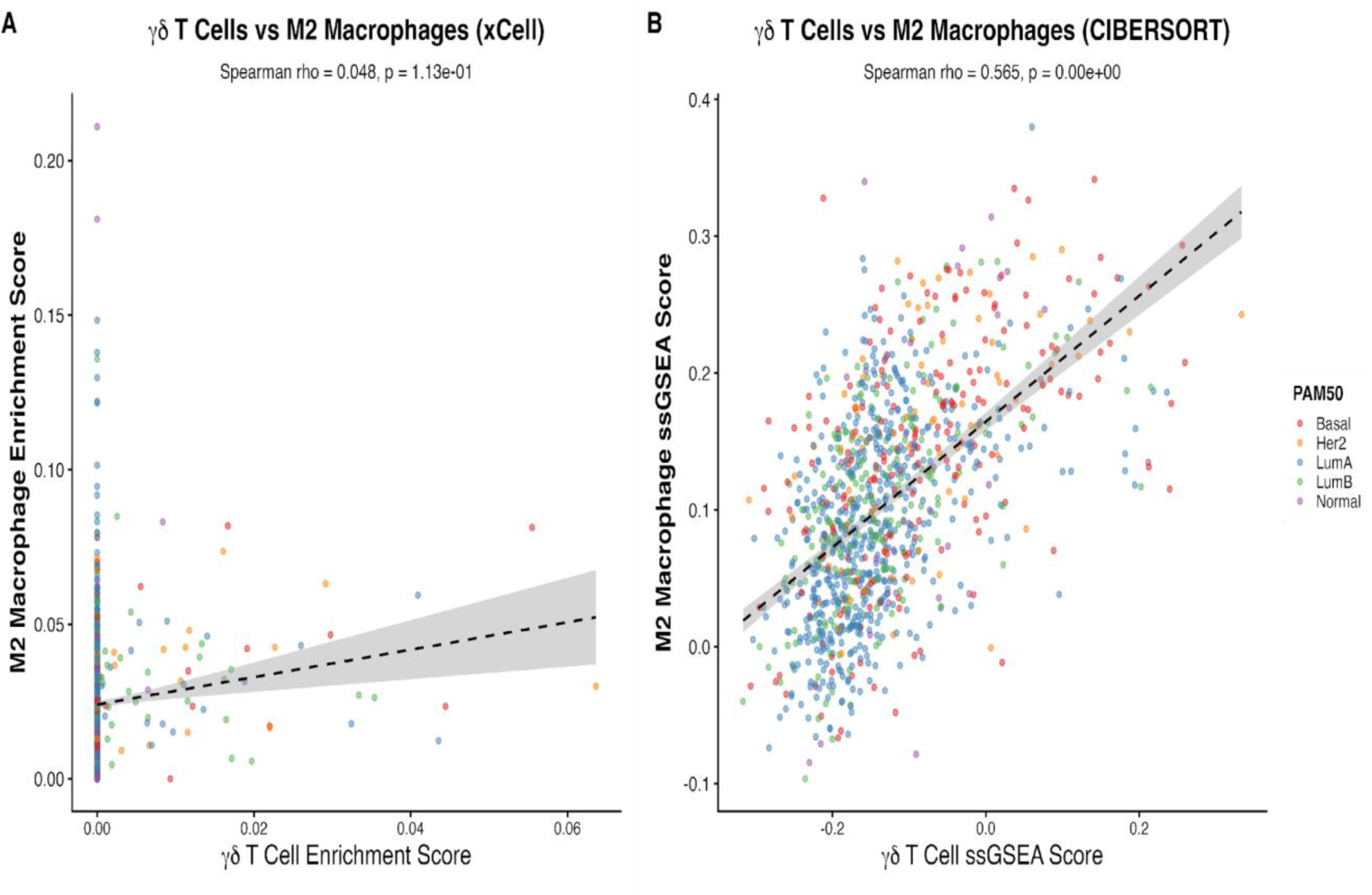
γδ T cell vs M2 macrophage enrichment. (A) xCell: ρ = 0.048, p = 0.11. (B) CIBERSORT-ssGSEA: ρ = 0.565, R² ≈ 0.32, p < 2.2 × 10⁻¹⁶. Points colored by PAM50 subtype.

The co-enrichment of γδ T cells and M2 macrophages has a clear biological basis. IL-17-producing γδ T cells promote M2 macrophage polarization through NF-κB signaling (12). Petroni and colleagues showed that in HR+ breast cancer, CDK4/6 inhibitor therapy triggers recruitment of IL-17A-secreting γδ T cells that reprogram TAMs into a CX3CR1+ immunosuppressive state (4). Our data suggest the circuit is not just therapy-induced but present at baseline implying that the γδ–M2 axis seems contributing to immune exclusion in ER+ disease even before treatment starts, and its disruption might help sensitize these tumors to immunotherapy. The strongest subtype-specific γδ–M2 correlation was in luminal B tumors (ρ = 0.59), which is clinically significant: these patients have worse outcomes than luminal A and are increasingly treated with CDK4/6 inhibitors. If they already have an elevated baseline γδ–M2 co-enrichment, therapy-induced amplification of this axis is a plausible resistance mechanism. Strategies that disrupt the circuit, whether through IL-17 pathway blockade, macrophage reprogramming with Class IIa HDAC inhibitors or CSF1R antagonists (18), or therapeutic γδ T cell modulation (19), deserve further investigation in combination regimens. Together, the two methods give a clearer answer than either alone would: CIBERSORT-ssGSEA uncovers the γδ–M2 signal, xCell confirms that it is a rare-cell signal that requires a sensitive scoring approach, and biology fills in why the co-enrichment makes sense.

### 2.5 Multivariate modelling reveals a two-component model of PD-L1

A multivariate linear regression predicting PD-L1 from immune cell scores and PAM50 subtype explained 49% of the variance (adjusted R² = 0.49). M1 macrophages, Tregs, and CD8+ T cells were the strongest positive predictors, consistent with PD-L1 being driven by both IFN-γ from an active cytotoxic response and the regulatory immune activity that typically accompanies it. PAM50 subtype contributed additional explanatory power beyond immune cell composition, with basal-like tumors retaining a high baseline PD-L1 that immune infiltration alone did not fully account for (Figure 5). This indicates to a two-component model of PD-L1 regulation. One component is immune-driven: M1 macrophages and CD8+ T cells secrete IFN-γ, which upregulates PD-L1 on tumor cells as an adaptive immune-evasion response. The other is tumor-intrinsic and tied to molecular subtype. Basal-like tumors have higher PD-L1 even after accounting for their higher immune infiltration, which suggests cell-autonomous regulation, possibly through oncogenic signaling pathways that directly activate the CD274 promoter. This distinction matters for biomarker development: measuring PD-L1 protein by immunohistochemistry cannot distinguish between these two sources, and a tumor with high PD-L1 from intrinsic activation may respond differently to checkpoint blockade than one whose PD-L1 is driven by active immune pressure. The 49% of variance captured by this two-component model also explains, at least in part, the subtype-specific correlation patterns seen earlier: where an immune response is present, PD-L1 tracks it; where it is absent but the subtype is basal-like, PD-L1 can still be elevated.

**Figure 5.**
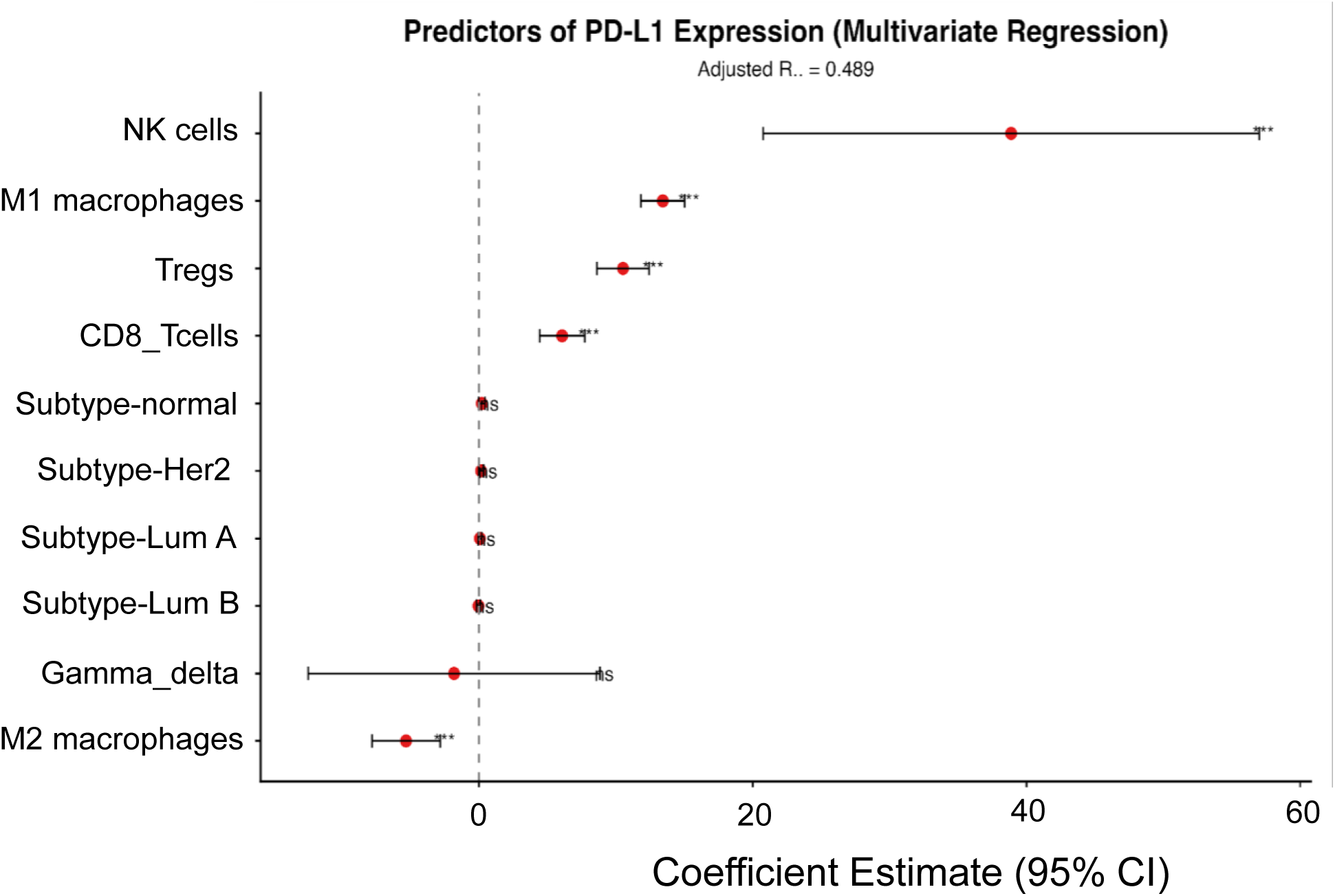
Forest plot showing multivariate regression coefficients predicting PD-L1 expression at 95% CIs and R² = 0.49.

### 2.6 Immune signaling context for the γδ–M2 axis

Expression pattern of genes implicated in the γδ–M2 axis was analyzed across the subtypes (Fig. S4), and was found to be varied in different subtypes. For instance, IFN-γ was highest in basal-like tumors as expected while IL-17A was uniformly low which is unsurprising given that only a small γδ T cell subset produces IL-17A, and the global transcriptomics may not be able to detect the changes occurring in small fraction of cell population. The CCL2, the chemokine implicated in γδ T cell recruitment (4), varied across subtypes with higher levels in basal-like and normal-like tumors while the expression of CX3CR1, the macrophage marker, of the immunosuppressive TAM varied and agreed to earlier observation (4). Although the global RNA seq analysis would not be able to detect the gene expression in cells producing low levels of any transcript, they are consistent with the pathway mechanisms that could drive the γδ–M2 co-enrichment, with CCL2 recruiting γδ T cells, those cells producing IL-17 (below bulk detection), and IL-17 reprogramming macrophages toward a CX3CR1+ M2 state. Confirming this sequence will require single-cell and spatial approaches, which would complete the pathway-level picture that the bulk analysis can only outline.

### 2.7 Immune composition adds modest prognostic information

The correlation between immune composition and patient survival was monitored across the subtypes. Results showed that the patients with high immune scores obtained using xCell, though individual immune populations had more complex prognostic relationships, tend to survive longer (Figure S5). The Cox model, incorporating PAM50 subtype and immune features, gave a moderate and expected concordance of 0.60. Overall survival in breast cancer depends heavily on stage, treatment, and molecular features that immune composition alone does not capture. When we interpreted these survival results, we observed that the TCGA cohort is not ideal for survival analysis because the follow-up times are uneven and treatment data are incomplete, and the cohort predates modern immunotherapy regimens. However, a concordance of 0.60 suggests that immune features add some prognostic information on top of subtype but not enough to build a standalone prognostic model.

### 2.8 Subtype-specific immune architectures at a glance

A heatmap of all immune cell estimates across 1,099 tumors were analyzed. Results showed that the subtype-specific architectures clearly (Figure 6A). Basal-like tumors clustered with high CD8+ T cell, M1 macrophage, and dendritic cell enrichment. Luminal tumors grouped together with elevated M2 macrophages, Tregs, and stromal signatures HER2-enriched and normal-like tumors fell in between. Composite microenvironment scores confirmed the subtype-level differences (Figure 6 B–D). Together, these panels summarize the central theme of the study: each breast cancer subtype operates under its own set of immune rules, and therapeutic strategies aimed at the tumor microenvironment must be designed with that subtype-level architecture in mind.

**Figure 6.**
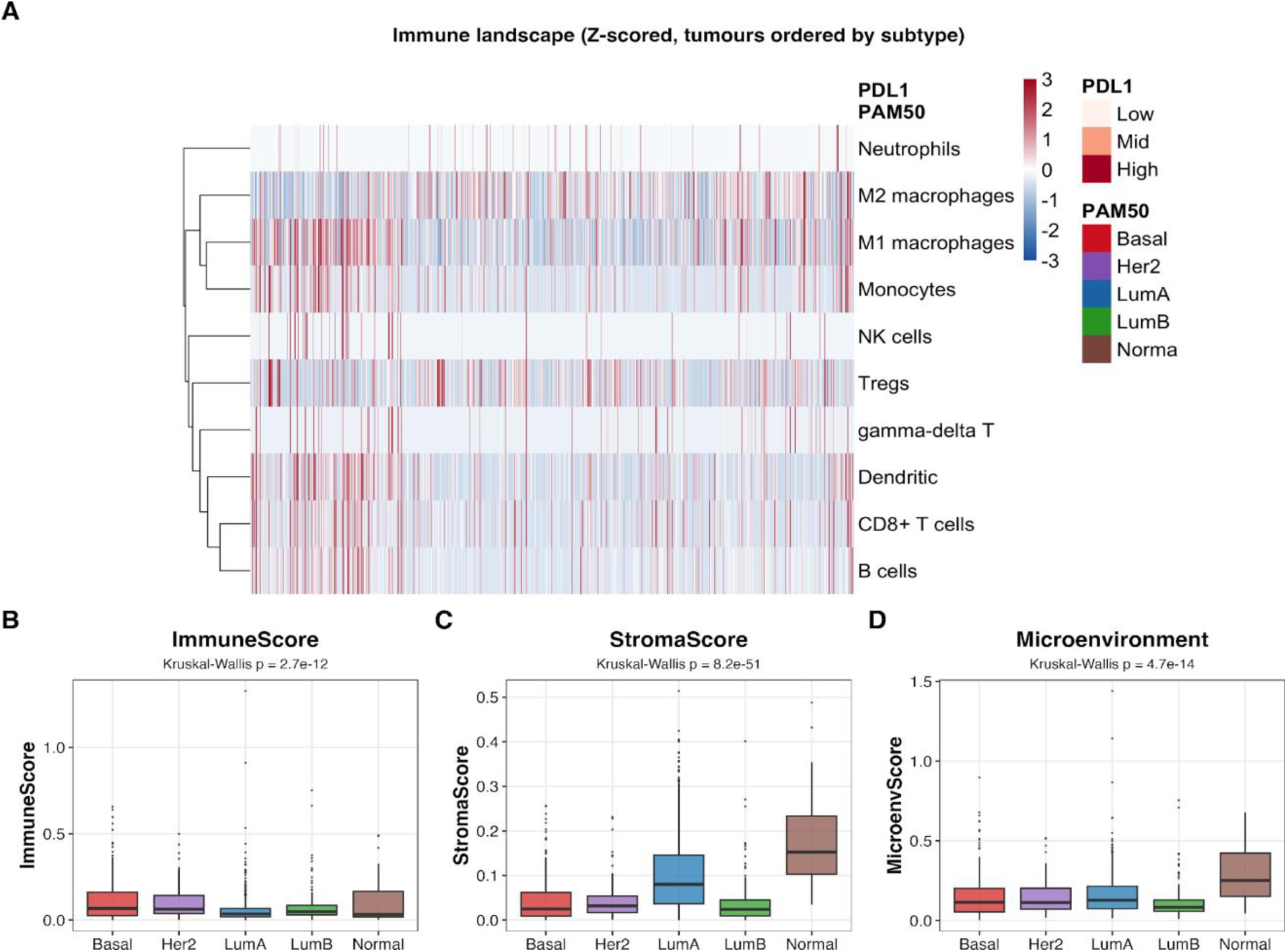
Subtype-specific immune architectures. (A) Heatmap of Z-scored immune cell enrichment across 1,099 tumors, ordered by PAM50 subtype; annotation bars indicate subtype and PD-L1 level (tertile). (B) Immune Score, (C) Stroma Score, and (D) Microenvironment Score by subtype.

In summary, earlier analysis of the breast cancer immune microenvironment using single deconvolution algorithm have produced inconsistent results limiting the usefulness of such information in cancer therapeutics. Here, we have brought forth the results obtained using two independent deconvolution methods on a same population of 1,099 breast tumors that could map the subtype-specific immune architectures and identified a γδ T cell–M2 macrophage co-enrichment pattern persisting across all PAM50 subtypes. For instance, CIBERSORT-ssGSEA detected this at a much higher confidence than xCell highlighting the significance of multi-method validation in characterizing rare immune populations. Although, the functional heterogeneity in the bulk RNA-seq data within the cell populations, the cytotoxicity from IL-17-producing γδ T cells, and the M2 macrophages, the LM22 signatures derived from purified blood immune cells may not perfectly capture tumor-adapted phenotypes, and the TCGA cohort being predominately of North American population are some of the factors that limits the use of this information for application, the available results from this study provide population-scale evidence for a baseline γδ–M2 immunosuppressive circuit, one that is present across all subtypes but may be especially relevant in ER+ disease and could be a candidate for therapeutic intervention. Further, a multivariate model accounting for immune composition and subtype explains nearly half of PD-L1 variance (adjusted R² = 0.49), reinforcing the view that PD-L1 regulation in breast cancer has both immune-driven and tumor-intrinsic components. Combined with the subtype-specific PD-L1 profiling, these findings may suggest for immunotherapy strategies designed around the specific immune wiring of each breast cancer subtype, rather than a one-size-fits-all approach.

## 3. Methods

### 3.1 Data acquisition and preprocessing

Gene expression data and clinical annotations for the TCGA-BRCA cohort were obtained through the TCGAbiolinks R/Bioconductor package (14). In brief, we retrieved STAR-aligned count files and TPM values for 1,231 samples. After restricting to primary tumors with available PAM50 classification, 1,099 cases were taken for further analysis: 197 basal-like, 82 HER2-enriched, 571 luminal A, 209 luminal B, and 40 normal-like. Raw counts were normalized to counts per million (CPM) and log2-transformed. Ensemble gene identifiers were mapped to HGNC symbols, retaining 19,938 protein-coding genes. Overall survival was defined as the number of days survived before death for deceased patients, and the number of days to the last follow-up after treatment for the surviving population.

### 3.2 Dual immune deconvolution

We chose two approaches deliberately. First, xCell, which scores 64 immune and stromal cell types using single-sample GSEA across 489 gene signatures, followed by spillover compensation to reduce inter-cell-type correlations (15). Second, a CIBERSORT-style ssGSEA in which we defined 22 immune cell type gene sets from the LM22 signature matrix (16), including curated γδ T cell markers (TRGC1, TRGC2, TRDC, TRDV2, TRGV9). Gaussian kernel estimation and normalization were carried out using the GSVA package (v2.4), and scores were computed as described in (17). Results obtained from both methods were cross-checked, giving higher confidence to features reported by both and flagging any that required closer examination.

### 3.3 Statistical analysis

Subtype differences were tested with Kruskal–Wallis tests and post-hoc Dunn tests (FDR-corrected). Associations between continuous variables used Spearman correlations, computed globally and within each subtype. Cross-method concordance was assessed by correlating matched xCell and CIBERSORT-ssGSEA estimates. Multivariate linear regression identified independent predictors of PD-L1, with the adjusted R² reported as a measure of overall fit. Survival was evaluated with Kaplan–Meier curves (log-rank test) and Cox proportional hazards models. All analyses used R (v4.5.1); p < 0.05 (two-sided) was considered significant.

## Acknowledgement

Authors are grateful to Anusandhan National Research Foundation (ANRF), Government of India for research grant vide reference No. CRG/2023/004799, and Prof. B. Jaya Kumar Singh and Dr Sayantani Ghosh, SDSOS, SVKM’s NMIMS Deemed to be University, for their critical comments on the manuscript.

## Conflict of Interests

Authors have not conflict of interest in the content of the paper. Authors have agreed for submission to PLOS ONE.

## Authors contributions

Diya Jain (DJ) is a graduate student of Bio-Medical Sciences, carried out this work under the supervision of Professor Hari S Misra (HSM). DJ conducted all the analysis, wrote draft paper and prepared figures. Hari S Misra is a Distinguished Professor of Biological Sciences, SVKM’s NMIMS Deemed to be University, Mumbai. HSM supervised the work carried out by DJ, wrote paper, data analysis and corresponded work for publication.

## Supplementary Materials

### Remarks

All supplementary figures were produced by the same analysis pipeline used for the main figures (scripts at code/01_prepare_data. R, code/02_deconvolution. R, code/03_analysis_and_figures. R). Source tables underlying each panel are provided as CSV files in the Tables/ directory of the project repository.

**Figure S1.**
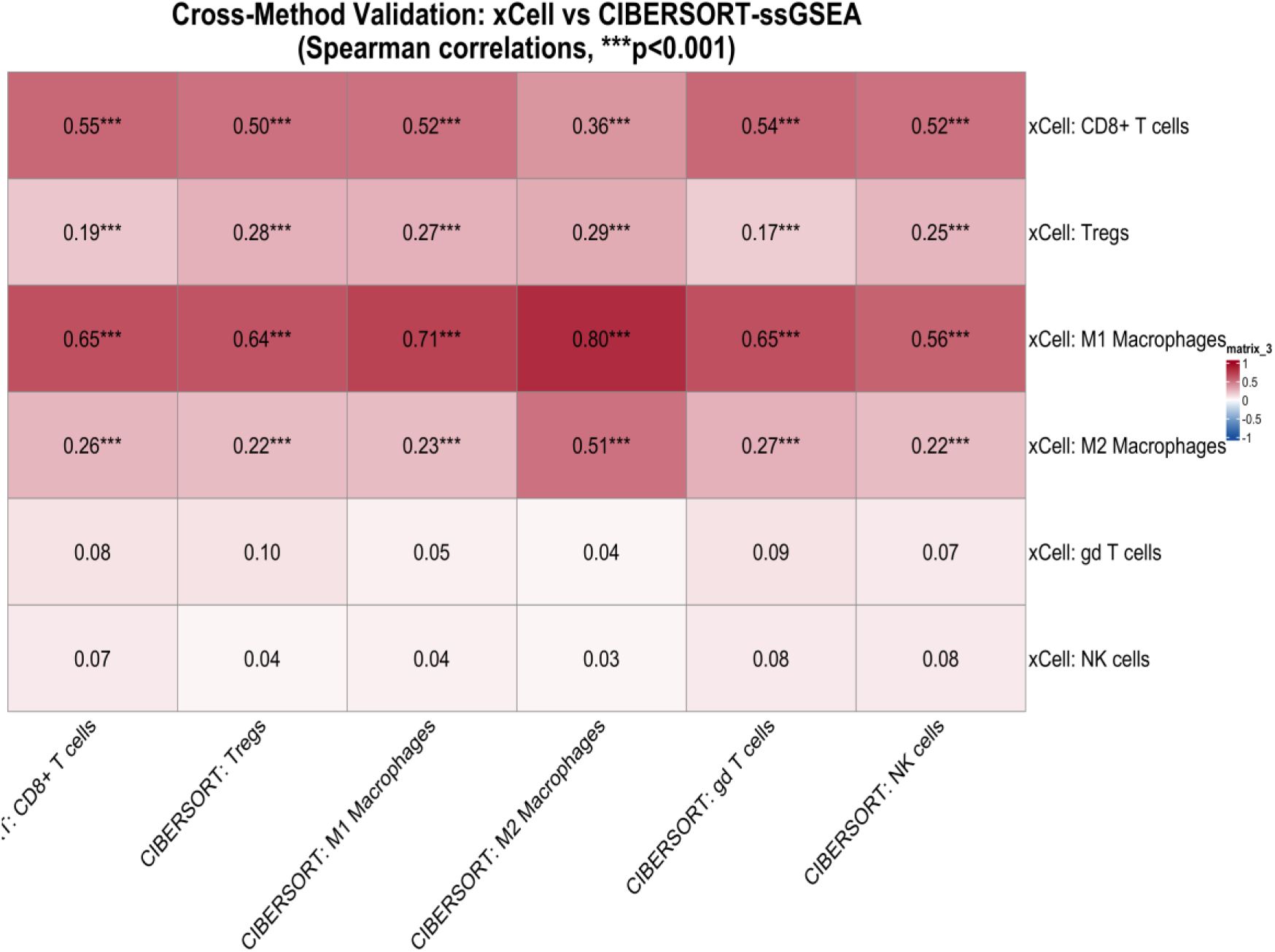
Spearman correlations between xCell scores and CIBERSORT-ssGSEA scores across matched cell types in all 1,099 TCGA-BRCA primary tumours. Strong concordance for CD8+ T cells, M1 macrophages, Tregs, and NK cells supports the validity of both pipelines. The γδ T cell – M2 macrophage axis is the one area where the methods diverge, and this divergence is addressed in the main text.

**Figure S2.**
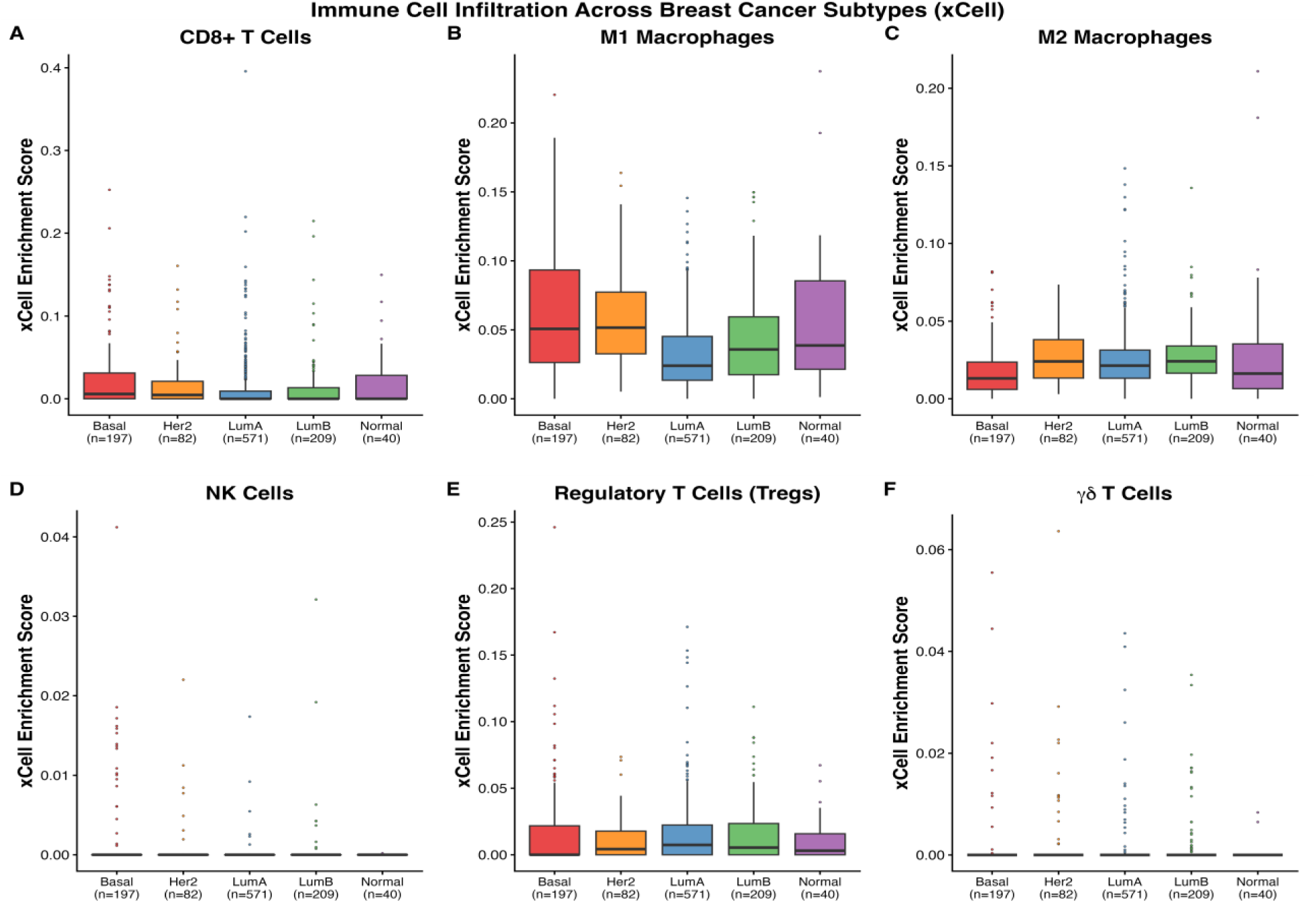
xCell enrichment scores across six immune populations. (CD8+ T cells (A), M1 macrophages (B), M2 macrophages (C), regulatory T cells (D), NK cells (E) and γδ T cells (F) for the five PAM50 subtypes (Basal n = 197, Her2 n = 82, LumA n = 571, LumB n = 209, Normal n = 40). Kruskal–Wallis p-values shown per panel. Condensed CD8 and M1 panels appear in Figure 2 of the main paper.

**Figure S3.**
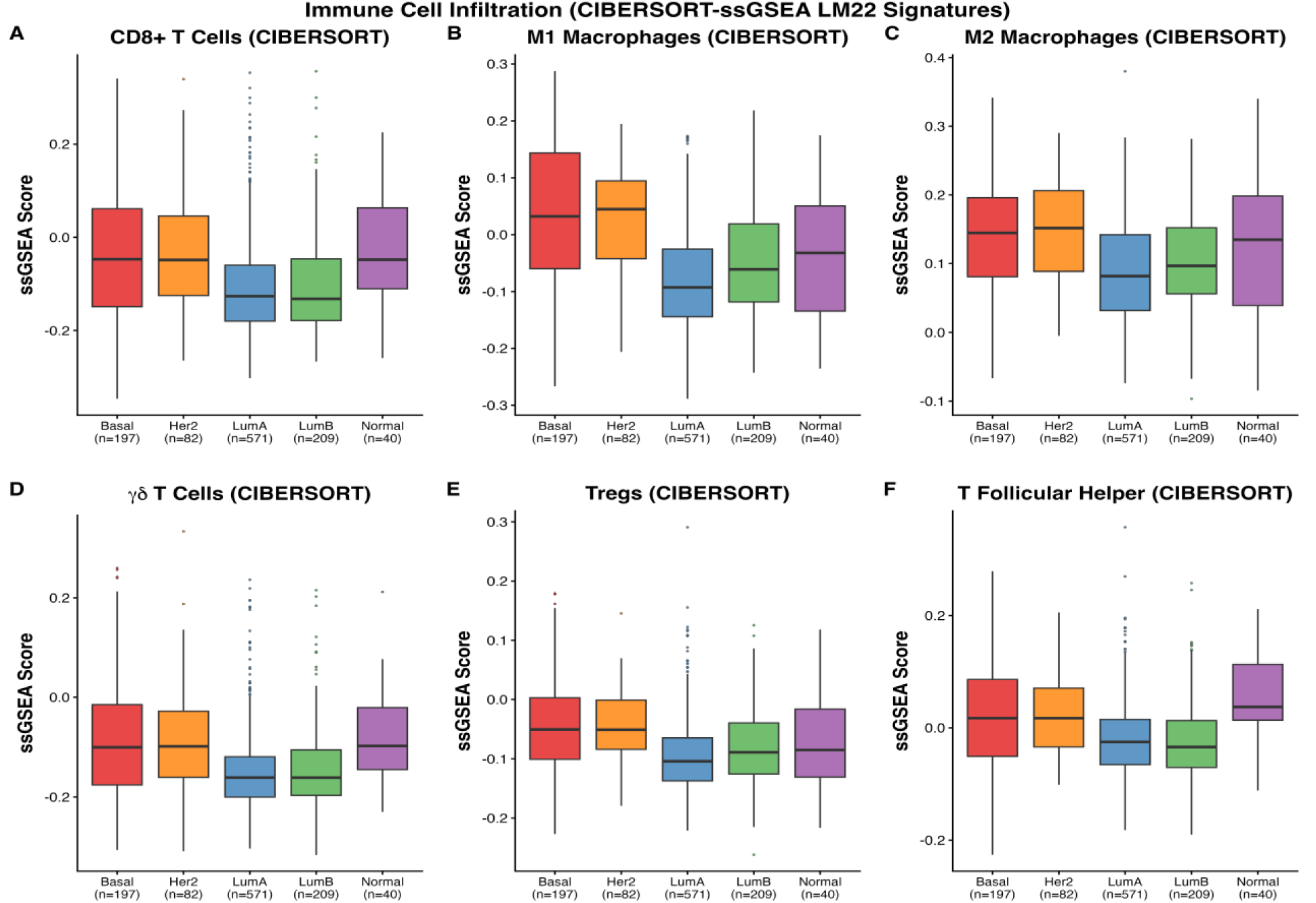
CIBERSORT-style ssGSEA scores using LM22 gene signatures across the same six immune populations and five PAM50 subtypes. The parallel panel layout lets readers compare method-to-method stratification directly. Condensed CD8 and M1 panels appear in Figure 2 of the main paper.

**Figure S4.**
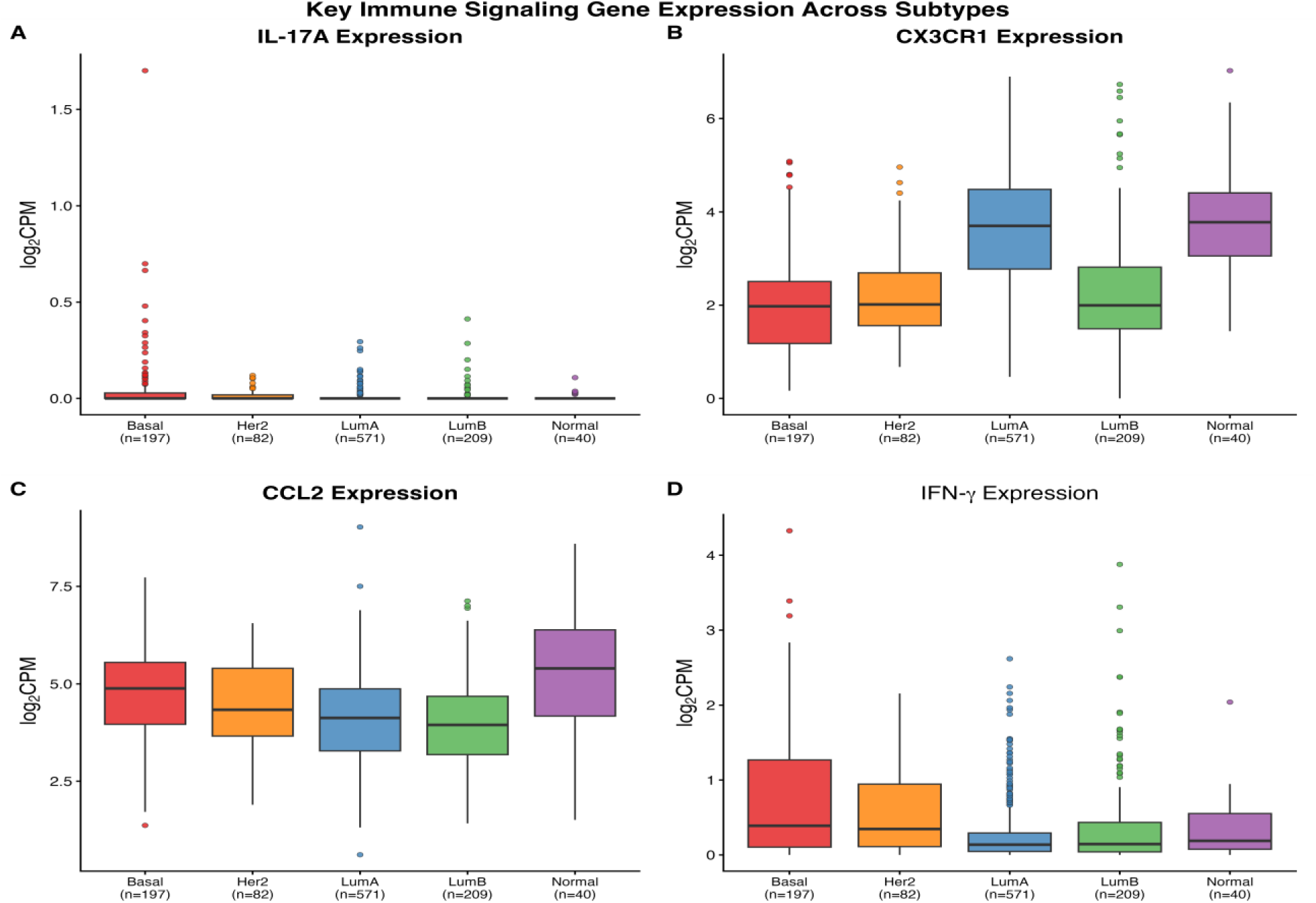
log2 CPM expression of IL-17A, CX3CR1, CCL2, and IFN-γ across PAM50 subtypes. These signalling molecules frame the cytokine context supporting the γδ T cell – M2 macrophage axis and help interpret why the axis surfaces in CIBERSORT-ssGSEA but not in xCell.

**Figure S5.**
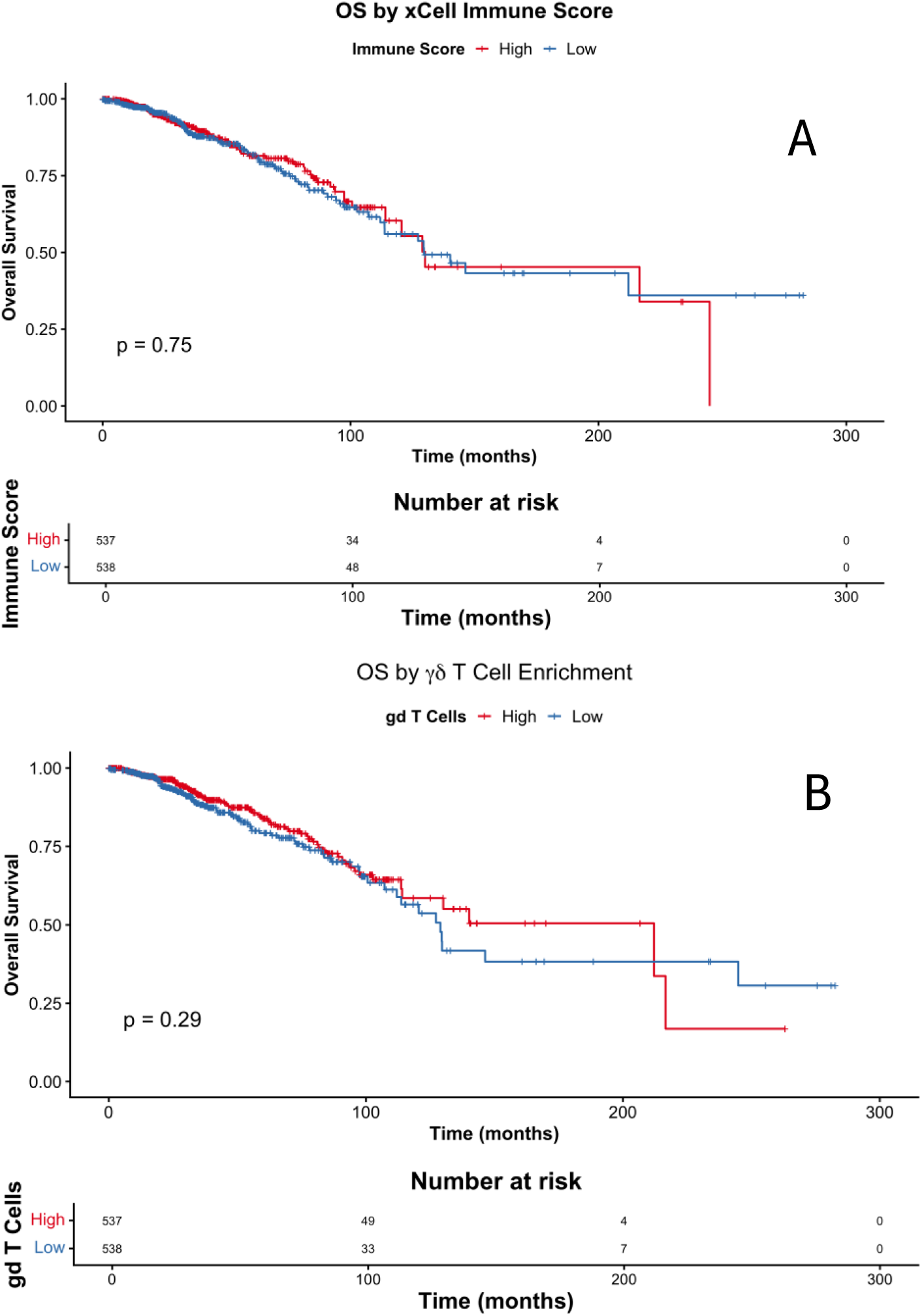
Overall survival stratified by tertiles of xCell and xCell γδ T cell Immune Scores. **A**. High-ImmuneScore tumours show a modest survival advantage (log-rank p reported in main Results). **B.** The curves echo the ImmuneScore trend and remain consistent with a protective effect of infiltrating lymphocytes.

## Notes

### Competing Interest Statement

The authors have declared no competing interest.

